# Task-specific effects of biological sex and sex hormones on object recognition memories in a 6-hydroxydopamine-lesion model of Parkinson’s disease in adult male and female rats

**DOI:** 10.1101/2022.02.15.480556

**Authors:** Claudia C. Pinizzotto, Aishwarya Patwardhan, Daniel Aldarondo, Mary F. Kritzer

## Abstract

Many patients with Parkinson’s disease (PD) experience impairments in cognition and memory with few therapeutic options currently available to mitigate them. This has fueled interest in determining how factors including biological sex and sex hormones might modulate higher order function in PD. Previous studies have investigated this in female rats and in gonadally intact and gonadectomized males, with and without hormone replacement, that received bilateral neostriatal 6-hydroxydopamine (6-OHDA) lesions to model PD. Barnes maze and What Where When Episodic-like memory testing showed that 6-OHDA lesions disrupted spatial working and episodic memory functions in both sexes, and that in males, androgen-sensitive behaviors could be rescued in subjects where circulating androgen levels were diminished. Here we tested similar animal groups using the Novel Object Preference (NOP) and Object-in-Place (OiP) tasks. This revealed two entirely different patterns of sex and sex hormone influence. First, for both tasks, 6-ODHA lesions impaired object discrimination in males but not females. Further, for the NOP task, 6-OHDA lesions disrupted discrimination in males rats independently of hormone status. And finally, 6-OHDA lesions impaired OiP performance in males regardless of whether androgen levels were high or low but had no effect on discrimination in gonadectomized rats given 17β-estradiol. Together with previous findings, these data identify the impacts of sex and sex hormones on cognition and memory in PD as behavioral task/behavioral domain specific. This specificity could explain why a cohesive clinical picture of endocrine impacts on higher order function in PD has remained elusive.

**HIGHLIGHTS:**

- 6-OHDA lesions impair Novel Object performance in male but not female rats.
- 6-OHDA lesions impair Object-in-Place performance in male but not female rats.
- Gonadectomy has no effect on 6-OHDA-induced deficits in Novel Object Preference.
- Estrogen replacement prevents 6-OHDA-induced Object-in-Place deficits in males.

## INTRODUCTION

Parkinson’s disease (PD) is the second most common neurodegenerative disorder[1–3]. In addition to motor deficits, e.g., bradykinesia, postural instability, resting tremor, many patients with PD also experience non-motor symptoms including impairments in cognition and memory[1, 2, 4–7]. These non-motor signs appear at or before the onset of motor symptoms in about one third of patients and have a cumulative prevalence of more than 80% overall[5–12]. Although the term ‘mild cognitive impairment’ distinguishes these deficits from those associated with Parkinson’s disease-related dementia (PDD), these disturbances are socially disabling, have a significant negative impact on patients’ quality of life, compromise their activities of daily living and increase caregiver burden[13–18]. Early occurrence of these impairments also predicts a more rapid clinical course of cognitive and motor decline, increased risk for freezing and falls and for developing PDD[9, 19, 20]. Finally, unlike the plurality of means available to treat motor symptoms, there are few therapeutic options that improve cognitive or memory function in PD[5, 10, 19, 20]. This has fueled interest in identifying and understanding the factors that influence cognitive and memory deficits in PD, including biological sex.

Many aspects of PD are strongly influenced by biological sex[21–27]. For example, males are nearly twice as likely as females to develop PD and to experience an accelerated increase in risk of illness with age[22, 28–30]. In addition, motor signs often present earlier and with greater severity in males, whereas females are more likely to experience fatigue, depression, gastrointestinal distress and excessive sweating[21, 22, 31, 32]. Although less well studied, sex differences have also been identified among the cognitive and memory impairments of PD[23, 31, 33]. For example, meta-analyses have identified male sex as conferring risk for cognitive and memory dysfunction in PD[34–36]. Clinical studies have also identified poorer spatial cognition, lower scores on tests of executive function, slower processing speed and reduced initiative in male compared to female patients as well as a greater likelihood for these deficits in males to progress from absent to mild and from mild to PDD [34, 37, 38]. These patterns suggest potentially important roles for circulating gonadal hormones in sex-specifically influencing cognition and memory in PD. Findings including increased risk of cognitive or memory disturbance in female PD patients with age[26, 39, 40] and of unusually low circulating testosterone levels in male PD patients[41, 42] suggest hypotheses of estrogen protection and/or of low levels of circulating testicular hormones as negatively impacting cognition and memory[27, 28]. However, there are inconsistencies in the clinical literature[39, 43, 44] including that examining potential benefits of hormone augmentation therapies for cognition and memory in PD[39, 43–47]. To resolve these open questions, this and other studies have turned to preclinical models to enable controlled investigation of the linkages between sex, sex hormones and the cognitive and memory disturbance associated with this condition.

Clinical attempts to identify the contributions of endogenous and exogenous gonadal steroid hormones to cognitive and memory disturbance in PD can be challenged by population variance in endocrine history, in hormone status, in key parameters of hormone replacement, e.g., timing, duration, formulations, and by a dearth of female patients. All of these variables can be controlled in animal studies. Moreover, the clear demonstrations of cognitive and memory deficits in preclinical rodent models of PD[48–51] make these especially well-suited for experimentally isolating sex and/or sex hormones impacts on disease-relevant higher order processes. The numbers of studies employing these means are growing and findings to date are promising. For example, studies in a genetic mouse strain that expresses the human A53T α-synuclein mutation[52] and in rats chronically treated with reserpine[53] both showed that object recognition memories were significantly more impaired in male compared to female subjects. In addition, studies in female rats in which 6-hydroxydopamine (6-OHDA)-induced nigrostriatal dopamine lesions were combined with ovariectomy found that dietary exposures to estrogen agonists or phytoestrogens (genistein, soy) improved spatial cognition in Morris water maze tasks[54, 55]. Previous studies in this laboratory have also assessed executive and episodic memory functions using Barnes maze testing[56] and the What Where When episodic-like memory (WWWhen ELM) task, respectively, in male rats where neostriatal 6-OHDA lesions were combined with gonadectomy (GDX) or with GDX paired with hormone replacement[57]. For both tasks, 6-OHDA-induced deficits in behaviors previously identified as androgen-sensitive were spared in lesioned rats where androgen levels were diminished. These and other data– including increased motor sensitivity to neurotoxic dopamine lesions in gonadally intact compared to GDX male rats[43, 58–60], suggest that the presence of androgens confers cognitive and mnemonic risk in Parkinsonian males. The objectives of this study were to determine whether similar rules applied to other domains of cognition or memory and to gain further inference about the neural networks where relevant, potentially protective hormone actions may be levied. Both objectives were met by assessing performance of separate cohorts of female, male and GDX male rats, with and without hormone replacement, that also received partial, bilateral neostriatal 6-OHDA lesions on the Novel Object Preference (NOP) or Object in Place (OiP) task. These tasks measure behaviors that are similar to the object recognition and visuospatial functions that are at risk in PD[61–65] and are relative free of potentially confounding effects of sex, sex hormones and dopamine systems on animals’ stress response, motor function and sensitivity to reward[66–69]. Further, the neural underpinnings engaged in these and other object recognition-based paradigms– including the WWWhen ELM task used previously, have been intensively studied[69–75]; this provides a fine-grained anatomical framework suitable for provisionally localizing key sites of biological sex and sex hormone actions. Finally, the necessary background information regarding hormone sensitivity is in hand for the NOP task[76]. However, because the effects of GDX and hormone replacement have never been evaluated for the OiP paradigm, the studies presented here include first assessments of OiP performance in non-lesioned female, male, GDX and GDX rats supplemented with estradiol or testosterone propionate.

## METHODS

### Animal Subjects

Subjects were adult male and female Sprague-Dawley rats (Taconic Farms, Germantown, New York, USA). Females ranged from 200-300 g and males ranged from 300-400g at the time of behavioral testing. All procedures involving rats were approved by the Institutional Animal Care and Use Committee at Stony Brook University and were performed in accordance with the U.S. Public Health Service Guide for Care and Use of Laboratory Animals to minimize their discomfort. Rats were pair-housed by sex and treatment group in standard translucent tub cages (Lab Products, Inc., Seaford, DE, USA) using ground corn cob bedding (Bed O’ Cobs, The Anderson Inc., Maumee, Ohio, USA) with food (Purina PMI Lab Diet: ProLab RMH 3000) and water available *ad libitum*. Cages and water bottles were made from bisphenol–free plastic (Zyfone) and rats were kept under a 12-h non-reversed light-dark cycle.

Animal groups included sham-lesioned females (FEM-SHAM, n = 13 for NOP, n = 10 for OiP) and female rats that received bilateral injections of 6-hydroxydopamine (FEM-OH, n = 9 for NOP, n = 10 for OiP). Male rats were distributed among 5 groups: males that were sham-lesioned (MALE-SHAM, n = 11 for NOP, n = 12 for OiP), gonadally intact males that were bilaterally injected with 6-OHDA (MALE-OH, n = 13 for NOP, n = 12 for OiP), and GDX, GDX TP and GDX E males that were bilaterally injected with 6-OHDA (GDX-OH, n = 10 for NOP, n =11 for OiP; GDXTP-OH, n = 6 for NOP, n = 12 for OiP, GDXE-OH, n = 9 for NOP, n = 12 for OiP). Additional cohorts of animals were also used to evaluate hormone effects in the OiP task. These included gonadally intact, unoperated female controls (FEM, n=12), gonadally intact, unoperated males (MALE, n = 12), males that were gonadectomized (GDX, n=11) and GDX males that were supplemented with testosterone propionate (GDXTP, n = 12) or 17β-estradiol (GDXE, n = 11).

### Surgeries

All surgeries were performed under aseptic conditions using inhalation of isoflurane (1% in oxygen) as anesthesia. Subcutaneous injection(s) of buprenorphine (0.03 mg/kg) or ketorolac (3 mg/kg) were given for postoperative analgesia.

#### Gonadectomy

Assigned male rats were gonadectomized 2 weeks prior to sham- or 6-OHDA injections, and 4 weeks before behavioral testing began. For this surgery a midline incision was made to the scrotum. The testes were then gently withdrawn, the vas deferens were ligated using silk sutures, and the testes were removed. For the animals assigned to the hormone replacement groups, slow-release pellets (Innovative Research of America, Sarasota, FL) were implanted into the surgical site. The pellets used were verified in previous studied to maintain plasma hormone levels within physiological limits, i.e., 3-4ng of testosterone propionate or 25pg of 17β-estradiol per millimeter [77–79]. Incision were then closed with surgical staples which were removed 5-7 days later.

#### Sham and 6-OHDA lesion surgeries

Partial bilateral neostriatal dopamine lesions were made by injecting 6-OHDA (6 μg/μL, Sigma-Aldrich, St. Louis, MO) dissolved in ascorbic saline (de-ionized water containing 0.9% NaCl and 0.1% ascorbic acid) using a glass micropipette. Thirty minutes prior to injection, rats received an intraperitoneal (IP) injection of desipramine hydrochloride (20 mg/kg, Sigma-Aldrich). For the surgery, anesthetized rats were placed in a stereotactic frame (Kopf Instruments, Tujunga, CA) and the skull was exposed using a midline incision. Next, burr holes were drilled at coordinates targeting the middle third of the rostral neostriatum in each hemisphere (AP: +0.5mm, ML: ±3.0mm, relative to Bregma). A glass pipette containing the 6-OHDA solution or vehicle secured to a Nanoject (Drummond Scientific, Broomall, PA) was then lowered through the burr holes to a depth of 5.8 mm below the brain surface. The pipette solution was then intermittently ejected (∼0.15 ul over 1 min) at 10-11 evenly spaced sites located 5.8 to 3.8 mm below the brain surface. This resulted in a total of ∼2ul of solution being injected per hemisphere. After the last ejection, the pipette was kept in place for an additional 10 min before being slowly withdrawn. Scalp incisions were closed using silk sutures and rats were given 2 weeks rest before behavioral testing began.

### Health and Hormone Monitoring

Rats’ weights and general health were monitored after GDX and/or 6-OHDA or SHAM lesion surgeries to assure that body mass progressively increased and that there were no signs of dehydration. The estrous cycles of the females were also tracked using intermittent vaginal lavage (saline) to collect cytological samples. These samples were collected on all days of behavioral testing. For males, the efficacies of hormone manipulations were assessed by dissecting and weighing the androgen-sensitive bulbospongiosus muscles at the time of euthanasia.

### Behavioral Testing

Habituation and behavioral testing took place during rats’ subjective night in a dedicated rat behavioral testing suite consisting of a central holding room and 5 adjoining sound-attenuated testing rooms. The testing room used was approximately 10 feet square, had even illumination and fixed, high contrast visual cues on three walls. Different testing arenas were used for the NOP and OiP tasks; both were placed on a table that was 36 inches high and located near the center of the room; a LogiTech webcam was suspended 2-3 feet above to record all sessions. The NOP arena was a translucent polypropylene rectangular enclosure that was 20 inches wide, 31 inches long and 12 inches deep. The OiP arena was a smaller, squarer shaped translucent polypropylene enclosure that was 15 inches wide, 21 inches long and 12 inches deep with a large fixed, high contrast panel marking one of its longer walls.

#### Testing procedures

Rats were habituated to handling and had undergone previous testing for novel open field and for either the What Where When episodic like memory or Barnes maze task. Additional task-specific habituation included a 1-hour acclimation to the central room of the testing suite prior to testing, and a 10 min session in the empty arenas.

##### NOP

Novel Object Preference testing consisted of a 3 min empty-arena trial, a 3 min Sample Trial and a 3 min Test Trial, all separated by 90 min inter-trial intervals spent in the home cage in the testing suite. During sample trials, pairs of identical objects were placed in adjacent corners along the short wall of the rectangular arena (4-inch clearance from all walls). To start the trial, rats were placed in the opposite end of the arena facing away from the objects. During test trials rats explored objects that were in the same positions, albeit one of which had been formerly sampled, and one of which was novel. The objects used were all approximately 3-4 inches tall, 3 inches across and cylindrically shaped; their distinguishing features were that one was smooth, metal and opaque and the other was glass and clear with shallow, vertically running ridges on its surface. The objects that served as sample vs. novel objects and their corner locations were counterbalanced across subjects. The arena and objects were cleaned with 70% EtOH before and after every trial.

##### OiP

Object-in-Place testing consisted of a 5 min Sample Trial and a 3 min Test Trial separated by a 5-minute inter-trial interval spent in the home cage in the testing suite. For both trials, rats were placed in an opaque start cylinder located at the center of the testing arena. After a 10 second delay, the cylinder was slowly lifted up and rats were free to explore 4 distinct objects placed near each of the arena’s four corners (4-inch clearance from all walls). The four objects used were similar in overall size and shape (5-7 inches tall, 2-3 inches across, cylindrical). However, one was made of aluminum and the other three were made of different grades of plastic. There were additional subtle differences in the objects’ contours and in their color/contrast (clear, bright, patterned and dark). During the test trials, rats explored the same four objects, albeit with two occupying original positions and two occupying positions that were switched relative to each other. The positions and pairs of objects that were switched vs. stationary were counterbalanced across subjects. The arena and objects were cleaned with 70% EtOH after each trial.

### Euthanasia

After behavioral testing was completed, rats were euthanized by transcardial perfusion. First, rats were deeply anesthetized by intraperitoneal injection of a ketamine (150mg/kg), xylazine (25mg/kg) solution. After deep reflexes were confirmed to be absent, rats were perfused with a flush of phosphate buffered saline (PBS, pH 7.4) followed by a 4% paraformaldehyde solution in 0.1M phosphate buffer (PB, pH 6.5, flow rate 30 ml/min for 5 min). This was then followed by perfusion using a 4% paraformaldehyde solution in 0.1M borate buffer (pH 9.5, flow rate 30 ml/min for 15 min). Following perfusion, the brains were removed and were post-fixed in 4% paraformaldehyde in 0.1M PB pH 7.4 for 24 hours and were then cryoprotected by immersion in 0.1M PB containing 30% sucrose. For the male rats, the bulbospongiosus muscle complex was also dissected out and weighed at this time.

### Tissue Processing and Histology

Cryoprotected brains were rapidly frozen in powdered dry ice and were serially sectioned (40 μm) on a freezing microtome. Immunohistochemistry for the dopamine-synthesizing enzyme tyrosine hydroxylase (TH) was performed on a 1 in 6 series of sections from sham- or 6-OHDA-lesioned animal as previously described[57] using an anti-TH mouse monoclonal antibody (LNC1 clone, EMD Millipore, Burlingame, MA, USA). Briefly, sections were rinsed in 0.1M PB, treated with 1% H_2_O_2_ in 0.1M PB, and incubated in heated sodium citrate buffer, pH 8.5 prior to a blocking step in tris-buffered saline (TBS), pH 7.4 containing 10% normal swine serum (NSS). Next sections were incubated for 4 days at 4 deg C in primary antibody, diluted 1:1000 in TBS containing 1% NSS. After buffer rinses, sections were transferred to a secondary antibody solution (biotinylated anti-mouse antibodies, Vector Laboratories, Burlingame, CA, USA) diluted 1:100 in TBS containing 1% NSS. After an overnight incubation (4 deg C), and following buffer rinses, sections were incubated for 5 hours (room temperature) in an avidin-complexed horseradish peroxidase solution (Vector Laboratories), prior to being rinsed and reacted using 3-3’diaminobenzidine (Sigma) as chromagen. Labeled sections were then slide-mounted in order, air dried overnight, dehydrated and placed under coverslips using Permount (Electron Microscopy Science, Hatfield PA, USA) prior to light microscopic evaluation.

### Data Analysis

#### Hormone Monitoring in Females and Hormone Manipulations in Males

Cytology samples obtained from vaginal lavage were used to track estrous cycle in female subjects. Samples were evaluated at low magnification using light microscopy and differential interference contrast illumination. Estrous cycle stages were identified as follows:

- Estrus-an abundance of cornified, anucleate epithelial cells.
- Diestrus-a predominance of leukocytes.
- Proestrus-a predominance of round, nucleated epithelial cells.

The effects of GDX and hormone replacement on circulating androgen levels were assessed by comparing weights of the dissected BSM muscle complex across male groups.

#### 6-OHDA Lesions

Digitized, low power brightfield images of TH-immunoreacted sections were captured using a Zeiss Axioskop light microscope outfitted with an Infinity 3 Lumenera digital camera. Images containing the neostriatum across its full rostro-caudal extent were archived for analysis. These images were imported into Fiji/ImageJ and the hand tool was used to outline the borders of the neostriatum in both hemispheres and to outline lesioned regions-if present-that were identified as zones where the intensity of TH immunostaining fell to background levels. Areal measures from both sets of outlines collected from all sections were used to obtain estimates of total neostriatal volume and total lesion volumes on per hemisphere bases; these values were used to quantify lesioned zones as percentages of total striatal volume. Total percent volumes (inclusive of both hemispheres) and ratiometric data representing lesion symmetry (hemispheric percent volumes; larger divided by smaller) were compared across animal groups and correlated within subjects to behavioral measures.

#### Behavior

Behavioral data were analyzed off-line from digitally recorded trials by trained observers who were blind to group using event-capture software (Behavioral Observation Research Interactive Software (BORIS) version 7.8.2, open access). All trials were analyzed for six major behaviors defined/described as:

> *Rearing*: total time (in seconds) rats spent standing on hind paws with fore paws either in contact with an object (without investigating), the arena walls or while free-standing.

> *Grooming*: total time (in seconds) rats spent preening (not differentiated by part of the body). *Ambulation*: total time (in seconds) rats spent engaged in walking/making forward motion with steps taken by all four paws.

> *Stationary Behavior*: total time (in seconds) rats remained sitting at one location and were not engaging in grooming, rearing or object investigation.

> *Latency to Investigate Objects*: time (in seconds) between the lifting away of the start cylinder and /or the start of a trial and rats’ first investigation of an (any) object.

> *Total Object investigation*: total time (in seconds) rats spent in contact with objects, using vibrissae or snout to actively investigate; forepaws may or may not also be in contact with the object.

Object investigation was further evaluated in task-specific ways as follows:

> *Sample Trial Spatial Bias*:

> NOP: total object investigation time (in seconds) spent with objects in each of the two arena corners.

> OiP: total object investigation time (in seconds) spent with objects in each of the four arena corners; total object investigation time spent with objects in arena’s top vs. bottom or left vs. right halves.

> *Sample Trial Object Bias:*

> NOP: total object exploration times (in seconds) spent with different sample objects, compared across subjects.

> OiP: total time (in seconds) spent with each of the four objects.

> *Test Trial Discrimination index:* assessed during Test Trials only. For female rats, these data were also stratified and compared according to estrous cycle/hormone status [relatively high (estrus, proestrus) vs. relatively low (diestrus)] on the day of testing].

> <u>NOP</u>: Total time (in seconds) rats spent investigating novel (NO) vs familiar objects (FO), expressed as percent of total object exploration time. This index was calculated by the following formula:

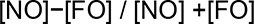

> <u>OiP</u>: Total time (in seconds) rats spent investigating objects in switched (Sw) compared to original (Or) positions, expressed as percent of total object exploration time. This index was calculated by the following formula:

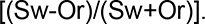

#### Statistics

All analyses were performed using IBM SPSS, Version 25 (SPSS, Inc., Chicago, IL, USA). The data were first assessed for descriptive statistics, including Levine’s F-test for equality of variance. The weights of the BSM complex, 6-OHDA lesion sizes and symmetries and discrimination indices (DI) were compared across groups using one-way analyses of variance (ANOVA). Allowed *post-hoc* testing used the Holm correction for multiple comparisons. Discrimination index data were additionally evaluated using within-groups one sample t-tests to determine whether DI values were significantly different from zero/chance. The relationships between DI and 6-OHDA lesion size and total object exploration times during Sample in individual subjects were assessed using Pearson’s regression analyses. All other comparisons, i.e., Major Behaviors, Object Bias, Spatial Bias, were evaluated using ANOVAs with repeated-measures designs. For these, Mauchly’s test for sphericity of the covariance matrix was applied and degrees of freedom were adjusted as indicated using the Huynh-Feldt epsilon. Where allowed, these comparisons were followed up by one-way ANOVAs to further identify significant group or object differences. Effect sizes were also assessed for all comparisons by calculating Cohen’s d (d) for t-tests or eta squared (*η*^2^) for all others. Finally, comparisons were made among male and female groups and among male groups with and without hormone manipulation to address a priori hypotheses about sex differences and about sex hormone sensitivities among the endpoints evaluated.

## RESULTS

### Hormone Manipulation and Monitoring

The weights of the androgen-sensitive bulbospongiosus muscle (BSM) complex served as a proxy for circulating androgen levels in male rats. The data in Table 1 show that the weights of this muscle group were larger in gonadally intact males (MALE, MALE-SHAM, MALE-OH) and in GDX rats supplemented with testosterone propionate (GDXTP, GDXTP-OH) compared to GDX rats (GDX, GDX-OH) and GDX rats given estradiol (GDXE, GDXE-OH). One-way ANOVAs that compared these data separately for animal used for NOP or OiP testing significant main effects of Group on BSM mass in all cases [NOP: F_(4, 41)_ = 61.18, *p* < 0.001, *η*^2^ =0.86; OiP: F_(3, 36)_ = 81.50, *p* < 0.001, *η*^2^ =0.87; OiP + 6-OHDA: *F*_(4, 44)_ = 87.52, *p* < 0.001, *η*^2^ =0.89]. Holm-corrected post-hoc comparisons further showed that for each of the studies, muscle weights of the GDX, GDX-OH, GDXE and GDXE-OH groups were similar to each other (NOP: *η*^2^ =0.09-0.101, OiP: *η*^2^ = 0.03; OIP + 6-OHDA: *η*^2^ =0.01-0.11) and were significantly lower than those in the MALE, MALE-OH, GDXTP and GDXTP-OH groups (*p* < 0.001-0.003, NOP: *η*^2^ =0.32-0.95; OiP: *η*^2^ = 0.6-0.98; OiP + 6-OHDA: *η*^2^ = 0.46-0.96).

**Table 1.** Group mean weights of the bulbospongiosus muscle (BSM) complex, expressed in grams ± standard error of the mean for male subjects tested on the Novel Object Preference (NOP) or Object in Place (OiP) tasks. Studies where comparisons included 6-hydroxydopamine (6-OHDA) lesioned subjects are designated + 6-OHDA. For all studies, BSM weights were significantly lower in rats that were gonadectomized (GDX) or GDX and given 17β estradiol (GDXE) compared to gonadally intact males and GDX rats given testosterone propionate (GDXTP, p< 0.001). This was true for groups that were unoperated, sham-operated, e.g., MALE SHAM, or that received partial bilateral neostriatal 6-OHDA lesions, e.g., GDX-OH).

| Table 1 | NOP + 6-OHDA | OiP | OiP + 6-OHDA |
| --- | --- | --- | --- |
| MALE | | 1.64± 0.04 g | |
| GDX | | 0.58± 0.02 g* | |
| GDXE | | 0.69± 0.04 g* | |
| GDXTP | | 1.12± 0.06 g | |
| MALE-SHAM | 1.58± 0.06 g | | 1.64± 0.03 g |
| MALE-OH | 1.45± 0.06 g | | 1.67± 0.07 g |
| GDX-OH | 0.57± 0.01 g* | | 0.59± 0.03 g* |
| GDXE-OH | 0.60± 0.02 g* | | 0.52± 0.03 g* |
| GDXTP-OH | 1.09± 0.14 g | | 0.96± 0.07 g |

Vaginal lavage and cytology performed during the weeks prior to and on the days of behavioral testing confirmed that all female rats were cycling regularly according to 4-day cycles. The data in Table 2 show the numbers of subjects by study that were in estrus (est), proestrus (pro) or diestrus (di) on the day of testing.

**Table 2.** Numbers of females (FEM), sham operated females (FEM-SHAM) or female rats that received partial bilateral 6-hydroxydopamine lesions FEM-OH) that were identified by vaginal cytology as having been in estrus (est), proestrus (pro) or diestrus (di) on the day of testing on the Novel Object Preference (NOP) or Object in Place (OiP) tasks. Studies where comparisons included 6-hydroxydopamine (6-OHDA) lesioned subjects are designated + 6-OHDA.

| Table 2 | NOP + 6-OHDA | OiP | OiP + 6-OHDA |
| --- | --- | --- | --- |
| FEM |  | 3 est; 4 pro; 5 di |  |
| FEM-SHAM | 4 est; 5 pro; 4 di |  | 3 est; 2 pro; 5 di |
| FEM-OH | 2 est; 3 pro; 4 di |  | 1 est; 5 pro; 4 di |

### 6-OHDA Lesions

Neostriatal lesions were not detected in sham-operated or unoperated rats. However, in all 6-OHDA-injected subjects, discrete, bilateral zones of sharply reduced TH-immunolabeling were evident (Fig 1E). Reconstructions of these zones revealed that lesion sites were roughly cylindrical in shape, were 0.5-1 mm in diameter, spanned most of the dorsoventral extent of the caudate nucleus and were centered near the midpoint of this nucleus at septal levels. Mean volumes subtended by these lesions were similar in all groups and accounted for no more than 24% of total, bi-hemispheric neostriatal volume. Measures of mean lesion symmetry (ratios of larger/smaller hemispheric lesion volumes) were also similar across groups and never exceeded a ratio of 2:1. Separate one-way ANOVAs (by behavioral study) confirmed that there were no significant main effects of Group on either lesion measure (*η*^2^ =0.07-0.17).

**Figure 1:**
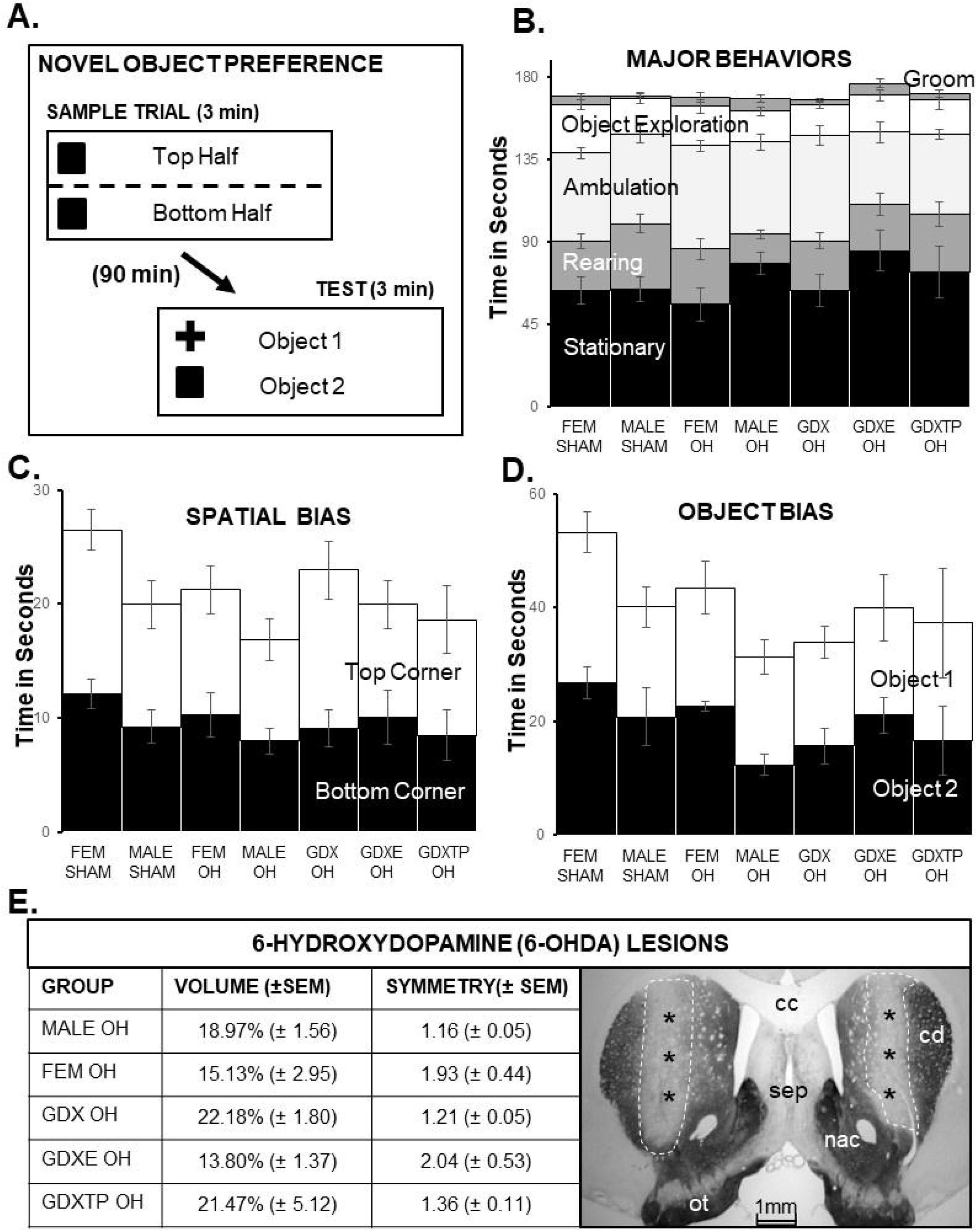
Sample trial data for the Novel Object Preference (NOP) task. (A) Schematic diagram showing trial structure for the NOP task. (B) Stacked bar graphs showing average amounts of time expressed as seconds ± standard error of the mean (SEM) that sham-operated female (FEM SHAM), sham-operated male (MALE SHAM) and six-hydroxydopamine (6-OHDA)-lesioned female (FEM OH), male (MALE OH), gonadectomized male (GDX OH) and GDX rats given 17-β estradiol (GDXE OH) or testosterone propionate (GDXTP OH) spent grooming (Groom, dark grey), exploring objects (white), ambulating (light grey), rearing (dark grey) or stationary (black) during the sample trial. Rats in all groups divided sample trial times similarly among these major behaviors. (C) Stacked bar graphs showing mean times expressed as seconds ± SEM that rats in each group spent exploring the object in the top (white) versus bottom half (black) of the arena during the NOP sample trial. No evidence of spatial bias was seen in any group. (D) Stacked bar graphs showing mean times expressed in seconds ± SEM that rats in all groups spent exploring sample objects when one of two object pairs was used. No evidence of object bias was seen in any group. (E) Table showing average volumes, expressed as percent total neostriatal volume and symmetry, expressed as larger over smaller hemisphere, ± SEM of 6-hydroxydopamine (6-OHDA) lesions for each lesioned group. There were no significant group differences for either measure. A representative low power photomicrograph is shown to the right. The outlined region of reduced tyrosine hydroxylase immunoreactivity marked by asterisks shows the lesion site in each hemisphere. Other abbreviations: cc-corpus callosum; sep-septal nuclei; nac-nucleus accumbens; ot-olfactory tubercle. Scale bar = 1mm.

### Novel Object Preference (NOP)

Animal subjects tested on the NOP task (Fig 1A) were gonadally intact females and gonadally intact males that were sham- or 6-OHDA lesioned; and GDX males and GDX rats supplemented with TP or E that were 6-OHDA lesioned. Across these groups, lesion sites subtended 14 to 22% of total neostriatal volume and lesion symmetry ratios ranged from 1.2 to 2 (Fig 1E).

### Sample Trials

#### Major Behaviors

During sample trials, rats in all groups spent most time stationary (∼70-100 sec), rearing (∼50-75 sec), or exploring objects (∼65-75 sec). Slightly less time was spent ambulating (∼35-50sec) and rats spent the least amounts of time grooming (∼8-16 sec) (Fig 1B). A repeated measures ANOVA confirmed that there were significant differences in amounts of times rats devoted to different behaviors [F _(2.33, 146.55)_ =132.84, p < 0.001, *η*^2^ = 0.68]. However, there were no significant main effects of Group and no significant interactions between Group and Behavior (*η*^2^ = 0.08 and 0.11, respectively).

#### Tests of Object Bias

Rats in all groups began investigating objects within a few seconds of the start of the trial; they also investigated each of the two sample items present roughly equally (Fig 1C). Within-groups, repeated-measures ANOVAs confirmed that there were no significant main effects of Object Position on exploration times for any of the groups. Effect sizes were also small (*η*^2^ = 0.001-0.043) with the exception of FEM-SHAM (*η*^2^ = 0.16) and GDX-OH rats (*η*^2^ = 0.25). Finally, within-groups, one way-ANOVAs that compared total object exploration times among subjects where sample objects were counterbalanced revealed no significant main effects of Object Type (Fig 1D) on this measure for any of the groups (*η*^2^ = 0.02-0.14).

### Test Trials

#### Major Behaviors

During test trials, rats in all groups spent most time stationary (∼70-100 sec) and more moderate times ambulating (∼35-50 secs), rearing (∼10-30 sec) or exploring objects (∼15-25 secs). The least amount of time was spent grooming (∼2-10 secs) (Fig 2A). A repeated measures ANOVA confirmed that times spent engaged in specific behaviors were significantly different [F_(2.75, 175.77)_ = 108.75, p < 0.001, *η* ^2^ = 0.63]. However, there were no significant main effects of Group and no significant interactions between Group and Behavior (*η*^2^ = 0.05 and 0.12, respectively).

**Figure 2:**
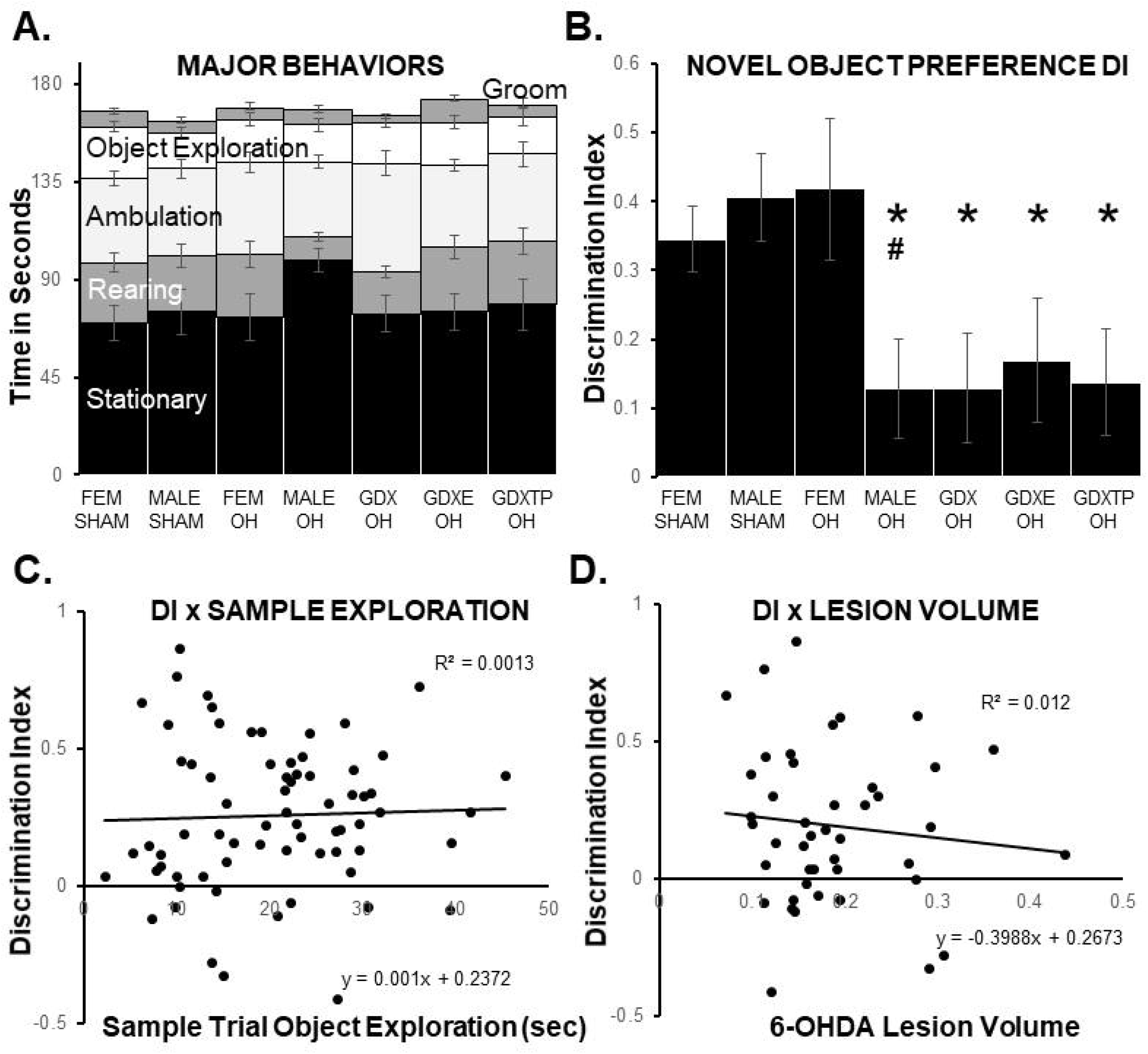
Test trial data for the Novel Object Preference (NOP) task. (A) Stacked bar graphs showing average amount of time in seconds ± (SEM) that sham-operated female (FEM SHAM), sham-operated male (MALE SHAM) and six-hydroxydopamine (6-OHDA)-lesioned female (FEM OH), male (MALE OH), gonadectomized male (GDX OH) and GDX rats given 17-β estradiol (GDXE OH) or testosterone propionate (GDXTP OH) spent grooming (Groom, dark grey), exploring objects (white), ambulating (light grey), rearing (dark grey) or stationary (black) during the NOP test trial. Rats in all groups divided test trial times similarly among these major behaviors. (B) Bar graphs showing group means ± SEM of calculated Discrimination Index (DI), i.e., ratios of times spent exploring novel compared to familiar objects, during the NOP test trial. Discrimination indices were significantly higher in FEM SHAM, FEM OH and MALE SHAM compared to MALE OH rats (#, p= 0.017-0.05), and DI in the MALE SHAM group was significantly higher than that of all other males (*, p= 0.013-0.05). Scatter plots comparing DI to total sample trial object exploration times (in seconds, C) and to total volumes of 6-OHDA lesions (expressed as total neostriatal volume, D). No significant correlations were found for either comparison. Trendlines, equations and R² values are shown.

#### Discrimination Index

Mean Discrimination index (DI) (Fig 2B) calculated for MALE-SHAM and FEM-SHAM rats were 0.41 and 0.34, respectively. The average DIs calculated for FEM-SHAM rats in estrus/proestrus was 0.37 and in diestrus was 0.30, respectively (*η*^2^ =0.037). Analyses of variance found no main effects of Sex (*η*^2^ = 0.026). For the FEM-OH rats, the average DI was 0.42; this was slightly but not significantly higher than the DI of the FEM-SHAM group. Within this group, mean DI for rats tested during estrus/proestrus was 0.55 and was 0.26 for rats tested during diestrus (*η*^2^ = 0.24). In comparison to lesioned females, the average DI calculated for MALE-OH rats was much lower (0.13). A one-way ANOVA comparing DIs across all four groups (FEM-SHAM, MALE-SHAM, FEM-OH, MALE-OH) identified significant main effects of Group [F_(3,42)_ = 3.85, p = 0.016, *η*^2^ = 0.22] and Holm-corrected post-hoc comparisons confirmed that DIs in the MALE OH rats were significantly lower than those of all other groups (FEM-SHAM, MALE-SHAM, FEM-OH, p = 0.017-0.05, *η*^2^ = 0.21-0.27). One-sample t-tests also showed that the DI’s for the FEM-SHAM, MALE-SHAM and FEM-OH group were all significantly different (greater) than zero/chance (p < 0.001-0.002; *d* = 0.17, 0.21 and 0.31, respectively).

Among 6-OHDA lesioned males, DIs were low in all three GDX groups (GDX-OH =0.13, GDXTP-OH = 0.14, GDXE-OH=0.17) (Fig 2B). These values were all comparable to those identified for MALE-OH rats (0.13) and were noticeably lower than DIs for the MALE-SHAM group (0.41). Analyses of variance that compared these DI valued confirmed that there were significant main effects of Male Group on DI ([F_(4,44)_ = 2.62, p = 0.048, *η*^2^ = 0.19]. Holm-corrected post hoc comparisons further showed that average DIs for the MALE-OH, GDX-OH, GDXE-OH and GDXTP-OH groups were all significantly lower than those of MALE-SHAM rats (p = 0.013-0.05, *η*^2^ = 0.21-0.31). Effect sizes for all non-significant comparisons were small (*η*^2^ = 0-0.006). A series of one-sample t-tests further showed that only the DI’s for the MALE-SHAM group were significantly different (greater) than zero/chance (p < 0.001, *d* = 0.21). Finally, correlations (Pearson’s) between individual DI scores and Total Object Exploration times measured during sample trials (Fig 2C) and between DI and measures of neostriatal lesion volume (Fig 2D) and symmetry (not shown) were all non-significant (sample exploration, r^2^ = 0.0013; lesion volume, r^2^ = 0.012; lesion symmetry, r^2^ = 0.0078).

#### Object-in-Place (Hormone Effects)

Animal subjects initially tested on the OiP task (Fig 3A) were gonadally intact females, gonadally intact males, GDX males and GDX males supplemented with TP or E.

**Figure 3:**
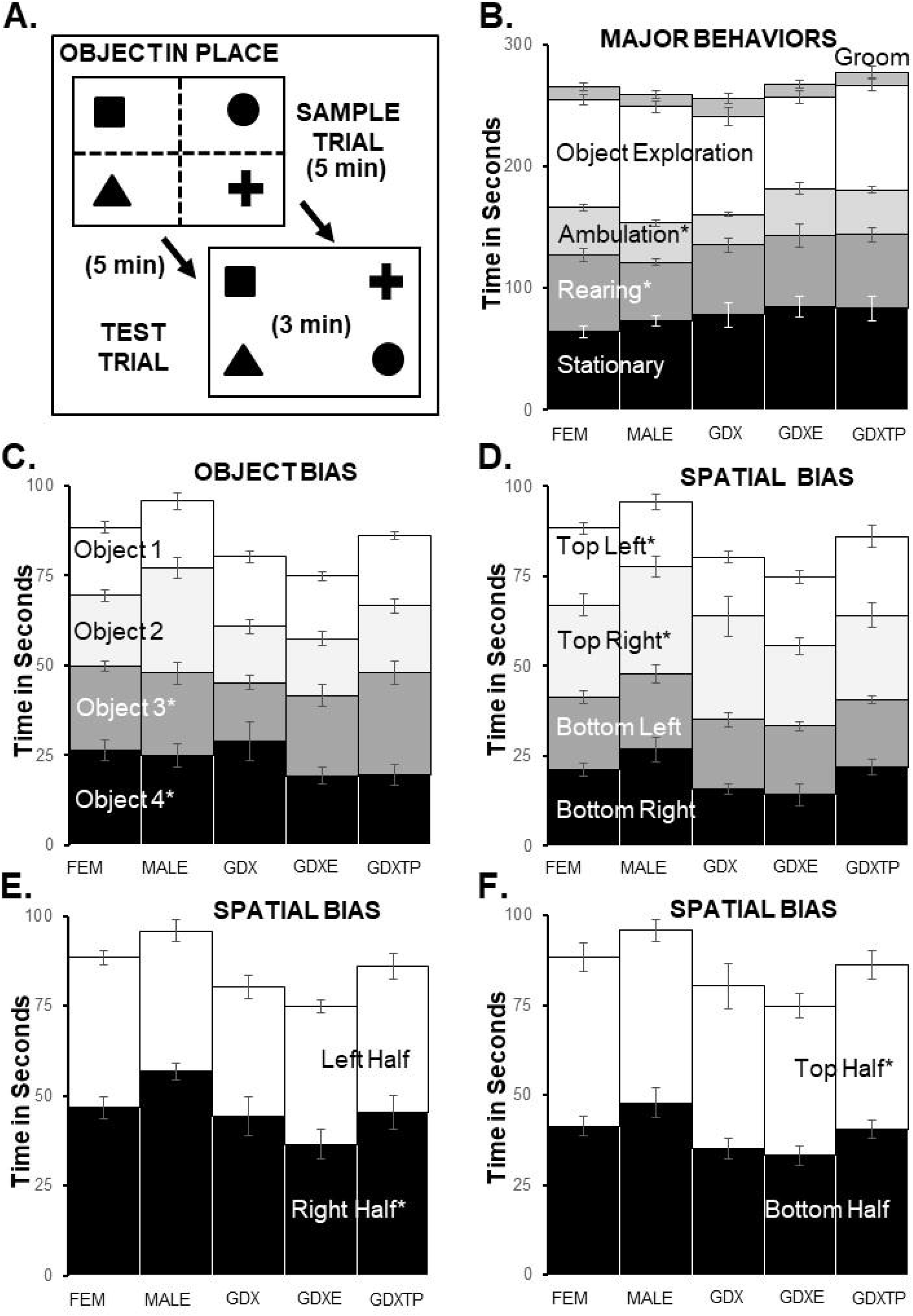
Sample trial data for the Object in Place (OIP) task. (A) Schematic diagram showing trial structure for the OIP task. (B) Stacked bar graphs showing average amounts of time expressed as seconds ± standard error of the mean (SEM) that female (FEM), male (MALE), gonadectomized male (GDX) and GDX rats given 17-β estradiol (GDXE) or testosterone propionate (GDXTP) spent grooming (Groom, dark grey), exploring objects (white), ambulating (light grey), rearing (dark grey) or stationary (black) during the sample trial. Most groups divided sample trial times similarly among these behaviors. However, FEM rats reared(*) significantly more than MALE rats (p= 0.025) and GDXTP rats spent significantly more time ambulating(*) than GDX rats (p= 0.002). (C) Stacked bar graphs showing group mean times in seconds ± SEM rats spent exploring each of four distinct sample objects (white, light grey, dark grey, black). Female rats spent significantly more time with Object 4 (* black) compared to Object 1 (white). GDX spent significantly more time with Object 4 (* black) than Objects 2 (light grey) or 3 (dark grey), while GDXTP rats spent significantly more time with Object 3 (* dark grey) compared to Objects 1 (white) or 2 (light grey) (p= 0.038-0.044). (D) Stacked bar graphs showing group mean times in seconds ± SEM rats spent exploring objects located in each of the arenas four corners (white, light grey, dark grey, black). MALE rats spent significantly less time in the Top Left (* white) compared to the Top Right (light grey) or Bottom Right (black) corners, and significantly more time in the Top Right (* light grey) compared to the Bottom Left (dark grey, p= 0.007-0.02) corner. Stacked bar graphs showing mean group times in seconds ± SEM rats spent exploring sample objects in the left (white) versus right half (black) of the arena (E) or in the top half (white) versus the bottom half (black) of the arena (F). MALE rats spent significantly more time with objects in the right compared to left half of the arena (* p< 0.001) and GDXE rats spent significantly more time exploring sample objects in top compared to the bottom half of the arena (* p= 0.015).

### Sample Trials

#### Major Behaviors

During sample trials, rats in all groups spent most time exploring objects (75-95 sec) or remaining stationary (∼65-85 sec), less time rearing (∼50-60 sec) or ambulating (∼25-40 sec) and little time grooming (∼10-15 sec) (Fig 3B). A repeated measures ANOVA identified significant differences in the amounts of time rats devoted to these behaviors [Behavior, F _(2.76,146.17)_ =201.38, p < 0.001, *η*^2^ = 0.79] and significant main effects of Group [F_(4.53)_ =4.65, p =0.003, *η*^2^ = 0.26]. However, no significant interactions between Behavior and Group were found (*η*^2^ = 0.08). Holm-corrected one-way ANOVAs that compared discrete behavioral data across MALE and FEM groups identified significant main effects of Sex (MALE vs. FEM) only for Rearing (F _(1, 22)_ = 5.75, p = 0.025, *η*^2^ = 0.21); these effects were driven by FEM rats rearing for an average of roughly 15 seconds more than MALE during the trial. For non-significant comparisons effect sizes (*η*^2^) ranged from 0.002-0.14. Similar comparisons among male groups identified significant main effects of Male Group (MALE vs. GDX vs. GDXE vs. GDXTP) only for Ambulation [F _(3,42)_ = 3.43, p = 0.025, *η*^2^ = 0.20] which was driven by GDXTP rats ambulating for an average of roughly 12 seconds more than GDX rats. For non-significant comparisons effect sizes (*η*^2^) ranged from 0.002-0.16.

#### Tests of Object Bias

Assessments of object exploration by object type showed that rats in all groups investigated each of the four items roughly equally (Fig 3C). A series of within-groups, repeated-measures ANOVAs confirmed that there were only three cases where main effects of Individual Object were significant [FEM: F_(3, 33)_ = 3.03, p = 0.043, *η*^2^ = 0.22; GDX: F_(1.42, 14.19)_ = 4.38, p = 0.044, *η*^2^ = 0.31; GDXTP: F_(2.01, 22.14)_ = 3.78, p = 0.038, *η*^2^ = 0.26). Female rats spent ∼8 seconds more with one object compared to all others. For GDX and GDXTP, main effects were driven by rats spending between 9-13 seconds more exploring one object compared to two others. For non-significant comparisons, effect sizes (*η*^2^) ranged from 0.15-0.17. Assessments of exploration by object location showed that rats in all groups spent similar amounts of time with objects located in each of the arena’s corners (Fig 3D). Within-groups, repeated-measures ANOVAs identified only one case where main effects of Object Location were significant [MALE: F_(1.95, 21.42)_ = 3.64, p = 0.045, *η*^2^ = 0.31]. These effects were driven by corner-to-corner differences in exploration of no more than 12 seconds. Rats also divided exploration times approximately equally for objects located in the arena’s top vs. bottom (Fig 3E) and left vs. right halves (Fig 3F). Within-groups, repeated-measures ANOVAs identified only two cased where differences in exploration were significant [GDXE: F_(1, 10)_ = 8.55, p = 0.015, *η*^2^ = 0.46; MALE: F_(1, 11)_ = 41.88, p < 0.001, *η*^2^ = 0.79]. The proportions of exploration times driving these effects were 55/45 and 60/40, respectively. For non-significant comparisons effect sizes(*η*^2^) ranged from 0.000-0.29.

### Test Trials

#### Major Behaviors

During test trials, rats in all groups spent most time stationary (∼50-100 sec) less time exploring objects (15-38 sec), rearing (∼20-45 sec) or ambulating (∼17-22 sec) and little to no time grooming (∼10-15 sec) (Fig 4A). A repeated measures ANOVA identified significant main effects of amounts of time were devoted to discrete behaviors [F _(2.45, 129.87)_ = 108.67, p < 0.001, *η*^2^ = 0.67], a significant main effect of Group [F_(4,53)_ = 7.26, p < 0.001,*η*^2^ = 0.35] and a significant interaction between the two [F_(9.80,129.87)_ = 5.22, p < 0.001, *η*^2^ = 0.28]. Follow-up, within-behaviors comparisons found no significant main effects of Sex, but did identify significant main effects of Male Group for Stationary Behavior [F _(3,42)_ = 7.63, p < 0.001, *η*^2^ = 0.35] and Object Exploration [F_(3,42)_ = 8.55, p < 0.001, *η*^2^ = 0.38]. Holm-corrected post hoc comparisons showed that these effects were driven by longer times that GDXE rats spent stationary compared to all other groups (p < 0.001-0.002, *η*^2^ = 0.36-0.52) and by shorter times that GDXE rats spent exploring objects compared to rats in all other male groups (p < 0.001-0.002, *η*^2^ = 0.38-0.46). For non-significant comparisons, effect size (*η*^2^) ranged from 0.01-0.20.

**Figure 4:**
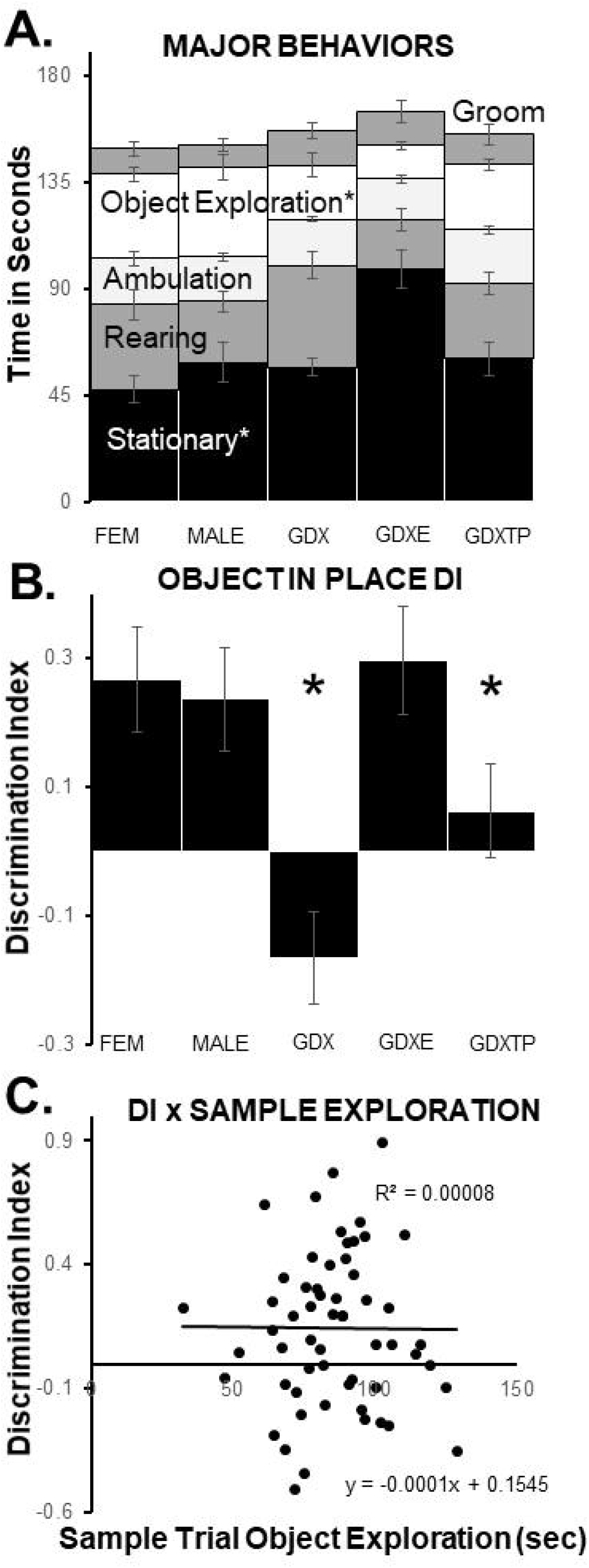
Test trial data for the OiP task. (A) Stacked bar graphs showing average amounts of time expressed as seconds ± standard error of the mean (SEM) that female (FEM), male (MALE), gonadectomized male (GDX) and GDX rats given 17-β estradiol (GDXE) or testosterone propionate (GDXTP) spent grooming (Groom, dark grey), exploring objects (white), ambulating (light grey), rearing (dark grey) or stationary (black) during the test trial. Most groups divided test trial times similarly among these behaviors. However, GDXE rats spent significantly more time stationary compared to all other groups (* p< 0.001-0.002) and significantly less time exploring objects compared to all other male groups (* p< 0.001-0.002). (B) Bar graphs showing group means ± SEM of calculated Discrimination Index (DI), i.e., ratios of times spent exploring moved compared to stationary objects during the OIP test trial. Discrimination indices were significantly lower in GDX and GDXTP rats compared to all other male groups (* p< 0.001). (C) Scatter plots comparing DI to total sample trial object exploration times (in seconds) show no significant correlations between these measures. Trendlines, equations and R² values are shown.

#### Discrimination Index

The average DI (Fig 4B) calculated for MALE rats was 0.21 and the corresponding value for FEM rats was slightly higher (0.27). There were minimal differences, however, among average DIs for female rats in estrus/proestrus (0.29) vs. diestrus (0.22) on the day of testing(*η*^2^ = 0.02). Analyses of variance that compared DI data across MALE and FEM subjects revealed no significant main effects of Sex on DI (*η*^2^ = 0.003) and one-sided t-tests confirmed that DI values in both groups were significantly different than zero/chance (p = 0.004-0.007, *d* = 0.28).

Among all male groups, average DIs (Fig 4B) were low for GDX (−0.17) and GDXTP (0.06) rats but were noticeably higher in the GDXE group (0.30). Analyses of variance that compared DI data across all male groups (MALE, GDX, GDXTP, GDXE) identified significant main effects of Male Group on DI [F_(3,42)_ = 7.00, p < 0.001, *η*^2^ = 0.33]. Holm-corrected post hoc comparisons further showed that DIs for the GDX group were significantly lower than those of MALE and GDXE rats (p ≤ 0.001, *η*^2^ = 0.40-0.47). The DI values for GDXTP rats, however, were not significantly different from either the lower DIs of GDX rats (*η*^2^ = 0.20) or the higher DI’s of the MALE and GDXE groups (*η*^2^ = 0.35 0.11 and 0.18, respectively). One-sided t-tests further showed that DI values for GDX rats were significantly lower than zero/chance (p = 0.021, *d* = 0.24); that DI values in the GDXTP group were not significantly different than zero (*d* =0.25); and that DI values in the MALE and GDXE groups were significantly higher than zero/chance (p = 0.003-0.007, *d* = 0.28). Correlations (Pearson’s) between DI and total Sample Trial object exploration times were not significant (r^2^ = 0.00008) (Fig 4C).

#### Object-in Place (6-OHDA Effects)

Animal subjects tested on the OiP task were gonadally intact females and males that were sham- or 6-OHDA lesioned; and GDX males and GDX rats supplemented with TP or E that were 6-OHDA lesioned. Across these groups, lesion sites subtended 15 to 24% of total neostriatal volume and lesion symmetry ratios ranged from 1.1 to 1.9 (Fig. 5A).

**Figure 5:**
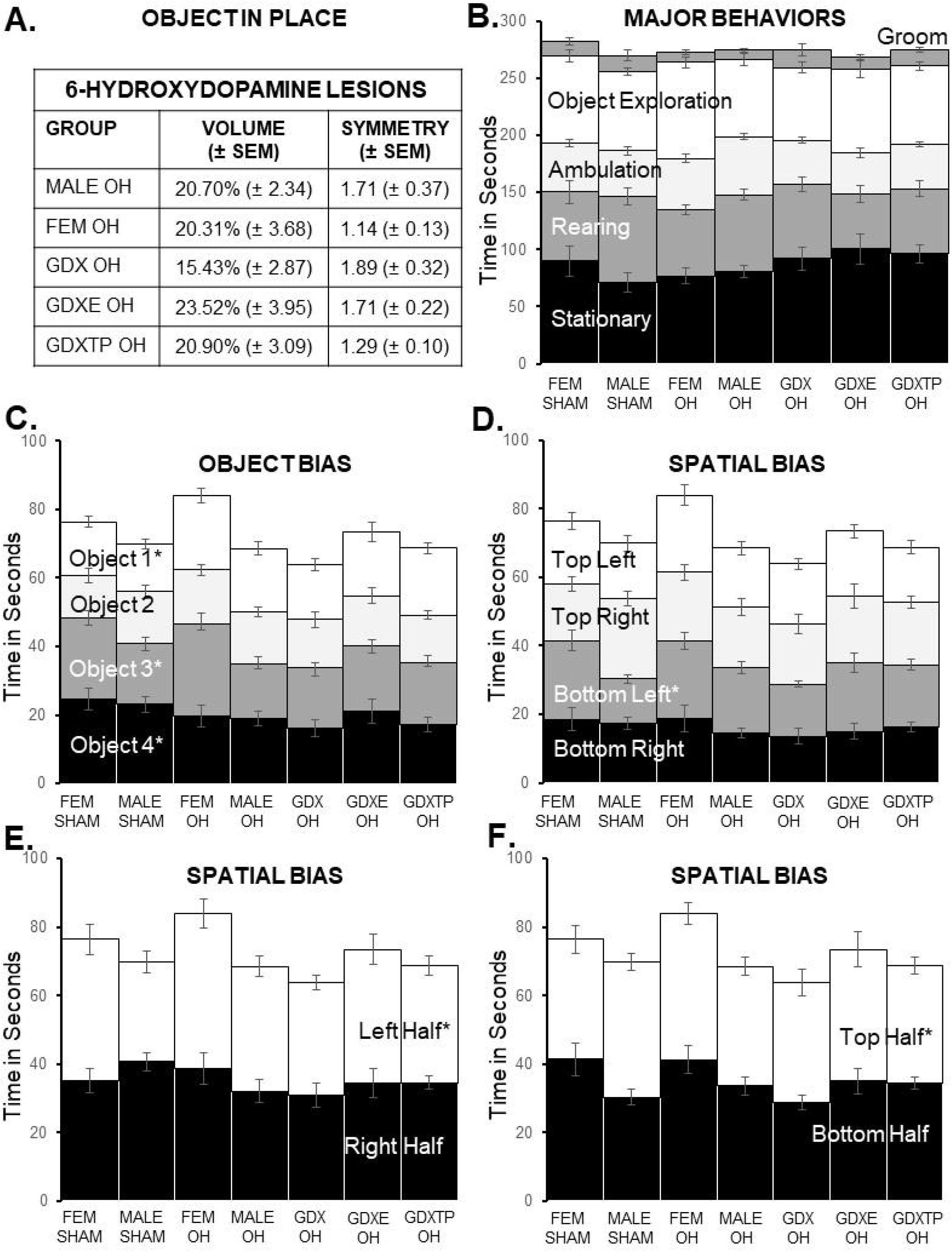
Sample trial data for the Object In Place (OIP) task. (A) Table showing average volume expressed as percent total neostriatal volume and symmetry, expressed as larger over smaller hemisphere, ± SEM of 6-hydroxydopamine (6-OHDA) lesions for each group. There were no significant group differences for either measure. (B) Stacked bar graphs showing average amounts of time expressed as seconds ± standard error of the mean (SEM) that sham-operated female (FEM SHAM), sham-operated male (MALE SHAM) and six-hydroxydopamine (6-OHDA)-lesioned female (FEM OH), male (MALE OH), gonadectomized male (GDX OH) and GDX rats given 17-β estradiol (GDXE OH) or testosterone propionate (GDXTP OH) spent grooming (Groom, dark grey), exploring objects (white), ambulating (light grey), rearing (dark grey) or stationary (black) during the sample trial. Rats in all groups divided sample trial times similarly among these major behaviors. (C) Stacked bar graphs showing group mean times in seconds ± SEM that rats spent exploring each of four distinct sample objects (white, light grey, dark grey, black). Sham operated female rats (FEM SHAM) spent significantly less time exploring Objects 1 (* white) and 3 (*dark grey) compared to the other two, and MALE SHAM rats spent significantly less time with Object 1 compared to Objects 3 and 4 (black), and significantly more time with Object 4 than 2 (light grey)(* p< 0.001-0.043). (D) Stacked bar graphs showing group mean times in seconds ± SEM rats spent exploring objects located in each of the arenas four corners (white, light grey, dark grey, black). MALE SHAM rats spent significantly less time with Objects in the Bottom Left (* dark grey) than in the Top (light grey) and Bottom Right (black) (p= 0.004-0.041). Stacked bar graphs showing mean group times in seconds ± SEM rats spent exploring sample objects in the left (white) versus right half (black) of the arena (E) or in the top half (white) versus the bottom half (black) of the arena (F). MALE SHAM rats spent significantly more time exploring objects in the right compared to the left half of the arena (E, * p= 0.035) and in the top compared to the bottom half of the arena (F, * p= 0.021).

### Sample Trials

#### Major Behaviors

During sample trials, rats in all groups spent most time stationary (∼70-100 sec), rearing (∼50-75 sec) or exploring objects (∼65-75 sec), less time ambulating (∼35-50 sec) and little time grooming (∼8-16 sec) (Fig 5B). A repeated measures ANOVA confirmed that there were significant differences in amounts of times rats devoted to discrete behaviors [F _(2.17, 154.27)_ =214.05, p < 0.001, *η*^2^ = 0.75]. However, there were no significant main effects of Group and no significant interactions between Group and Behavior (*η*^2^ = 0.09-0.11).

#### Tests of Object Bias

Rats in all groups investigated all four sample items approximately equally (Fig 5C). A series of within-groups, repeated-measures ANOVAs confirmed that main effects of Individual Object were only significant for the FEM-SHAM [F_(3, 27)_ = 7.26, p = 0.001, *η*^2^ = 0.45] and MALE-SHAM groups [F_(2.56, 25.60)_ = 4.20, p = 0.019, *η*^2^ = 0.30]. For FEM-SHAM rats, these effects were driven by rats spending ∼11 seconds less with two objects compared to the other two. In the MALE-SHAM group, effects were driven by rats spending ∼4-10 seconds longer with Object 1 compared to 3 and 4, and about ∼8 seconds more with Object 4 compared to Object 2. For non-significant comparisons, effect sizes (*η*^2^) ranged from 0.051-0.23. Rats also divided observation times roughly equally among objects located in each of the arena’s corners (Fig 5D) and in the arena’s left vs. right (Fig 5E) and top v. bottom (Fig 5F) halves. A series of within-groups, repeated-measures ANOVAs only identified significant main effects for the MALE-SHAM Group, for Object Corner, for Left/Right, and for Top/Bottom arena halves [F_(2.41, 24.10)_ = 5.17, p = 0.01, *η*^2^ = 0.34; F_(1,10)_ = 5.92, p = 0.035, *η*^2^ = 0.37; F_(1, 10)_ = 7.49, p = 0.021, *η*^2^ = 0.43, respectively]. Post hoc comparisons showed that in all cases, the effects were driven by MALE-SHAM rats spending ∼4-10 seconds less exploring one corner (Bottom Left) compared to all others. For non-significant findings, effect size (*η*^2^) ranged from 0.000-0.21.

### Test Trials

#### Major Behaviors

During test trials, rats in all groups spent most time stationary (∼60-80 sec), rearing (∼25-40 sec), exploring objects (∼20-32 sec) or ambulating (∼20-25 sec) and very little time grooming (∼5-12 sec) (Fig 6A). A repeated measures ANOVA confirmed that significantly different amounts of time were devoted to each behavior [F_(1.94, 137.65)_ = 214.13, p < 0.001, *η*^2^ = 0.75]. However, there were no significant main effects of Group and no significant interactions between Group and Behavior (*η*^2^ = 0.10 and 0.08, respectively).

**Figure 6:**
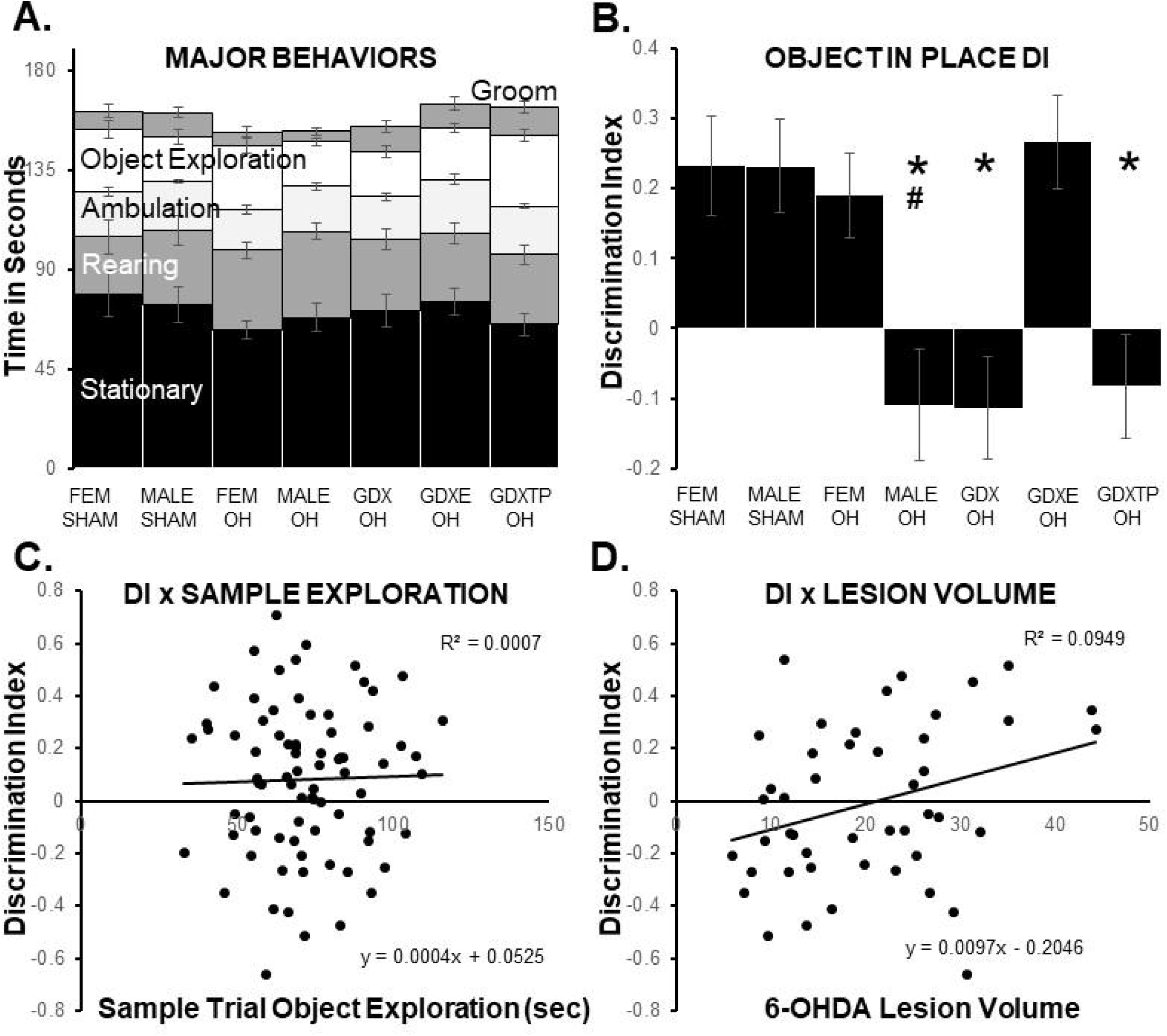
Test trial data for the Object In Place (OiP) task. (A) Stacked bar graphs showing average amounts of time expressed as seconds ± standard error of the mean (SEM) that sham-operated female (FEM SHAM), sham-operated male (MALE SHAM) and six-hydroxydopamine (6-OHDA)-lesioned female (FEM OH), male (MALE OH), gonadectomized male (GDX OH) and GDX rats given 17-β estradiol (GDXE OH) or testosterone propionate (GDXTP OH) spent grooming (Groom, dark grey), exploring objects (white), ambulating (light grey), rearing (dark grey) or stationary (black) during the test trial. Rats in all groups divided test trial times similarly among these major behaviors. (B) Bar graphs showing group means ± SEM of calculated Discrimination Index (DI), i.e., ratios of times spent exploring moved compared to stationary objects during the OiP test trial. Discrimination indices were significantly higher in FEM SHAM, FEM OH and MALE SHAM compared to MALE OH rats (#, p= 0.004-0.009) and DI in the MALE SHAM and GDXE OH groups were significantly higher compared to all other male groups (*, p< 0.001-0.05). Scatter plots hydroxydopamine (6-OHDA) lesions (expressed as total neostriatal volume (D). A significant positive correlation was found between DI and 6-OHDA lesion volume [D, r(76) = 0.31, p= 0.037)] but no significant correlations were found for sample object exploration. Trendlines, equations and R² values are shown.

#### Discrimination Index

Mean DIs (Fig 6B) calculated for MALE-SHAM and FEM-SHAM rats were both 0.23. There were also minimal differences in the average DIs calculated for FEM-SHAM rats that were in estrus/proestrus (0.22) vs. diestrus (0.25) on the day of testing (*η*^2^ =0.006). Analyses of variance revealed no significant main effects of Sex on DI (*η*^2^ = 0.00). The average DI for FEM-OH rats was 0.19; this was slightly but not significantly lower than the mean DI of the FEM-SHAM group (*η*^2^ = 0.01). The mean DI’s for FEM-OH tested during estrus/proestrus was 0.16 and was 0.23 for FEM-OH rats tested during diestrus (*η*^2^ = 0.035). The average DI calculated for the MALE-OH group was −0.11. This was markedly lower than the DIs of MALE-SHAM rats of both female groups. A one-way ANOVA comparing DIs across these four (FEM-SHAM, MALE-SHAM, FEM-OH, MALE-OH) identified significant main effects of Group [F_(3,39)_ = 5.80, p = 0.002, *η*^2^ = 0.31]. Holm-corrected post-hoc comparisons confirmed that DIs in MALE-OH rats were significantly different (lower) than all other groups (p = 0.004-0.009, *η*^2^ = 0.30-0.34). One-sample t-tests further showed that only the DI’s for the FEM-SHAM, MALE-SHAM, and FEM-OH were significantly different (greater) than zero/chance (p = 0.003-0.005; *d* = 0.19-0.23).

Among all males, DIs were lowest in the GDX OH (−0.11 and GDXTP-OH (−0.08) groups (Fig 6B). For GDXE-OH rats, the average DI was higher (0.27) and similar to that of MALE-SHAM rats (0.23). Analyses of variance that compared DI data across these groups identified significant main effects of Male Group on DI ([F_(4,53)_ = 7.08, p < 0.001, *η*^2^ = 0.48]. Holm-corrected post hoc comparisons further showed that average DIs for the GDX-OH, MALE-OH and GDXTP-OH groups were all significantly lower than those of the MALE-SHAM and GDXE-OH groups (p <0.001-0.05, *η*^2^ = 0.32-0.38) and that the DIs of GDXE-OH and MALE-SHAM rats. One-sample t-tests also showed that the DI’s of MALE-SHAM and GDXE-OH groups were significantly different (greater) than zero/chance (p = 0.003 and 0.001, respectively, d = 0.22 and 0.23, respectively and that the DI for the GDXTP-OH group was significantly different/less than zero/chance (p = 0.001; *d*= 0.26). Finally, correlations (Pearson’s) between individual measures DI and Total Object Exploration times measured during the Sample Trials (Fig 6C) and between 6-OHDA lesion symmetry (not shown) were not significant (sample exploration, r^2^ = 0.0007; lesion symmetry, r^2^ = 0.0004). However, a significant positive correlation was found between DI and 6-OHDA lesion volume [r(76) = 0.31, p = 0.037)] (Fig 6D).

## DISCUSSION

Meta analyses identify male sex as a risk factor for cognitive and memory dysfunction in PD[34–36]. Male patients are also more likely to develop cognitive and memory deficits early, to experience a more rapid course of cognitive decline and to be at greater risk for cognitive and memory deficits to progress to PDD[34, 37, 38]. These sex-specific features raise critical questions about the impacts of gonadal steroid hormones on cognition and memory in PD. Our lab has approached these questions using preclinical models of early, premotor stages of PD. Specifically, we have behaviorally tested rats where partial bilateral, neostriatal 6-OHDA lesions were paired with hormone monitoring in adult females and with GDX and hormone replacement in adult males using Barnes maze[56] and WWWhen ELM [57] tasks. Both studies showed that 6-OHDA lesions impaired performance in males and in females and that in males, androgen-sensitive behaviors were selectively spared from lesion-induced deficits in subjects where circulating androgen levels were decreased[56, 57]. Here we examined the performance of similar groups of animals tested for Novel Object Preference (NOP) or for Object-in-Place (OiP) recognition memory. These analyses brought to light new patterns of sex and sex hormone influence. First, unlike outcomes from Barnes maze or WWWhen testing, for both the NOP and OiP tasks 6-ODHA lesions significantly impaired discrimination in males but not females. Further, GDX and hormone replacement in 6-OHDA lesioned males affected NOP and OiP performance differently than previously seen and differently from each other. For the NOP task, where discrimination is impaired by GDX in an androgen-sensitive and estrogen-insensitive manner[76], 6-OHDA lesions disrupted recognition memory to similar degrees in all male subjects independently of their hormone status. However, for the OiP task, shown here to be disrupted by GDX in an estrogen-sensitive and androgen-insensitive manner, 6-OHDA lesions impaired performance in GDX-OH and GDXTP-OH rats but not in the GDXE-OH group. The behavioral task/behavioral domain-specific effects of sex and sex hormones on cognition and memory seen in this PD model are discussed further below, along with working hypotheses about their possible origins.

### Isolating Impacts on Cognition and Memory

Cognitive and memory impairments in PD patients include deficits in different forms of object recognition memory and visuospatial function[61–63]. This study extended earlier preclinical investigation of the effects of biological sex and sex hormones on cognition and memory to behaviors that closely match these at-risk operations using well-validated, widely used single trial object recognition tasks. As in earlier studies, partial bilateral neostriatal 6-OHDA DA lesions were used to model early, pre-motor stages of PD in male and female rats, and this PD model was paired with estrus cycle monitoring in females, and with GDX with and without estradiol or testosterone propionate replacement, in males. The use of animal models mitigates a number of the challenges faced in clinical studies of PD. However, experimental variables of sex, sex hormones and neostriatal dopamine lesions, as well as certain elements of the tasks themselves, bring challenges of their own for confidently ascribing group differences to impacts on cognition and/or memory. For example, studies in rats and mice have identified significant sex differences and sex hormone effects on experimentally induced nigrostriatal dopamine lesions– including those produced by neostriatal 6-OHDA injections[23, 80–85]. Although these effects are largely relegated to smaller-sized lesions, consensus findings show that injections of the same volumes of 6-OHDA produce larger DA lesions in males compared to females; in gonadally intact compared to GDX males; and in GDX males supplemented with dihydrotestosterone compared to GDX males given estradiol[23]. The generation of systematically smaller 6-OHDA lesions in androgen-intact males here could account for the observed sparing of NOP and OiP discrimination in 6-OHDA-lesioned female compared to male rats, and for the preservation of OiP discrimination in GDXE-OH compared to all other 6-OHDA lesioned male groups. Importantly, however, no significant group differences were found among measures of lesion volume or hemispheric lesion symmetry. Further, no significant negative correlations were observed between either lesion metric and NOP or OiP performance. Because bilateral neostriatal 6-OHDA lesions can also produce unwanted side effects of dysphagia and/or adipsia[86], it is also important to note that all animals were shown to have continually gained weight and maintained good hydration throughout the course of the studies.

The tasks chosen for analysis were selected first for their relevance to PD. Both the NOP and OiP paradigms measure discrete, well characterized behaviors that are similar to the simpler and more complex/integrative recognition memory deficits that have been noted in non-demented PD patients[38, 61, 62]. It is a further benefit that both tasks also measure object recognition memories in ways that are similar to human testing[69]. Moreover, the NOP and OiP tasks require habituation but no formal pre-training; rely on spontaneous rather than motivated behaviors; require minimal physical exertion; and produce minimal stress in the animal subjects[66, 67, 69]. Thus, these tasks operationally mitigate the well-known, potentially confounding impacts of biological sex, sex hormones and/or dopamine systems on rodents’ learning styles and speeds, sensitivity to reward, motor function and responses to stress[87–97]. Single trial object recognition-based tasks are also of proven utility and longstanding use in studying the effects of sex and sex hormone on higher order information processing[66]. As discussed below, the NOP task has also been used extensively to evaluate cognition and memory in preclinical rodent models of PD[98].

Although object recognition paradigms appear to be simple, their successful application requires that close attention be paid to potential confounds that can be introduced by the objects used and the spatial contexts in which they are presented[67, 70, 72]. For example, it is critical that sample and test objects have similar valence for the animal subjects and that the testing environment includes visible spatial cues. To guard against object bias, both studies used objects that shared general features of approximate size and general shape and were distinguished by more subtle characteristics, including material (glass, plastic, metal), color (high contrast, transparent) and surface texture (smooth, grooved, contoured). Measures of latency to initiate object investigation (any object) and total object exploration times were also tracked to assure that the objects used engaged similar levels of interest, encouraged approach and minimized neophobia. In addition, the objects used in sample vs. test trials were counterbalanced among rats within groups. To rule out contributions of object or spatial bias, detailed analyses were also made of rats’ object explorations by individual object type and by object location during sample trials. No systematic group differences in these measures were found. Largely separate cohorts of rats and different testing arenas were also used for the two paradigms to minimize potential confounds arising from test/re-test contingencies. And finally, it has been previously argued that the amounts of time rats spend on investigating sample objects has the potential to influence object memory tested at a later timepoint [99–102]. Because both paradigms used sample trials of fixed duration, measures of total sample trial object exploration were also evaluated as a function of DI in each animal subject to rule out sample exposure as a factor. These regression analyses revealed no significant correlations between the amount of time spent exploring sample objects and object discrimination measured in during the test trial. As an additional control, all subjects were evaluated for times spent on major ‘off-task’ behaviors including ambulation, rearing, grooming and remaining stationary. With very few exceptions rats in all groups apportioned times spent engaged in these discrete behaviors similarly in and across sample and test trials. The outcomes from these ancillary analyses sum to align group differences identified in test trial measures of object discrimination with the combined effects of sex, sex hormones and 6-OHDA lesioned subjects on intended target processes of object recognition memories.

### Sex Hormone Sensitivity of the OiP Task

A major objective of this study was to determine whether gonadal hormones influence object recognition memory functions in 6-OHDA lesioned male rats. Previous studies examining similar questions for other tasks relied on available *a priori* knowledge about the effects of GDX and hormone replacement on these processes in non-lesioned animals [56, 57, 103, 104]. For the NOP task, hormone sensitivity has been extensively studied in female rats and potentiating or protective effects of estrogens have been found among rats tested at differing stages of the estrous cycle and following ovariectomy and estradiol replacement[101, 105–109]. Fewer studies have examined the effects of gonadal hormones on NOP performance in male rodents[110, 111]. Among these are previous studies from our lab using the same procedures for GDX and the same procedures and apparatus for NOP testing used here. These studies showed that NOP discrimination was significantly impaired by GDX for both shorter (1.5 hrs.) and longer (4 hrs.) intertrial delays and that at both timepoints NOP discrimination deficits were attenuated in GDXTP but not GDXE rats[76]. While TP can act as an androgen and/or be metabolized to estradiol, the absence of appreciable rescue in the GDXE group identified NOP discrimination as an androgen-sensitive, estrogen-insensitive process[76]. Corresponding information about the impacts of GDX and hormone replacement for the OiP task, however, were not previously available. Thus, the present studies included an initial evaluation of OiP performance in gonadally intact MALE, GDX, GDXE and GDXTP male rats. These studies showed that OiP discrimination was significantly impaired by GDX and that DI deficits were attenuated in GDXE but not GDXTP rats. This indicates estrogen sensitivity and androgen-insensitivity, which is consistent with earlier studies in gonadally intact male rats showing that OiP performance was enhanced by local perirhinal cortical injections of estradiol or estrogen receptor agonists and was impaired by perirhinal infusions of estrogen receptor antagonists or aromatase inhibitors[112]. However, while the aromatase results implicate roles for estrogens derived from local testosterone metabolism in perirhinal cortex, the present study suggests that additional endogenous estrogens are also likely to contribute to the protection of OiP performance in GDX rats. In fact, the minimal sparing of OiP evident in the GDXTP group suggests that exogenous estrogen stimulation, i.e., estrogen acting at sites and/or concentrations that exceed the capacity of that derived from aromatase conversion, may play a major role in the behavioral rescue observed in the GDXE group.

### Previous Studies of Object Recognition Memory in Preclinical Rodent Models of PD

Cognitive and memory impairments in PD include deficits in object working memory, visual recognition memory and in integrated, visuospatial information processing[10, 38, 61, 62, 113]. The NOP paradigm has been successfully used to demonstrate similar deficits in an array of preclinical rodent models of PD[98]. For example, significantly impaired novel object discrimination has been reported in genetic rodent models of PD including mice with selective inactivation of mitochondrial transcription factor A in dopamine neurons (MitoPark)[114], mice that overexpress the human α-synuclein gene (Thy1-aSyn mice)[115], mice expressing the human A53T variant of α -synuclein[52] and rats where the PARK7 gene has been knocked out[116]. Deficits in the NOP task have also been identified in rats and mice where bilateral or unilateral nigrostriatal dopamine system neurodegeneration has been induced by intracerebral injections of MPTP, 6-OHDA or pesticides[117–122]; by intranasal administration of MPTP[123]; by exposure to environmental toxins including rotenone or paraquat[124–127]; by chronic treatment with reserpine[53, 128–132]; and by focal injections of either preformed fibrils or small soluble α-synuclein oligomers[133–136]. These data show that deficits in novel object recognition memory are highly robust to study-to-study differences in rodent species, in the means used to model PD pathophysiology and in testing regimens, including the use of identical vs. distinct sample items. However, there are a small number of studies where experimentally induced dopamine lesions had no reported effects on NOP performance. These include two studies where PD was modeled using bilateral neostriatal 6-OHDA similar to those used here[137, 138]. However, in both NOP was tested over unusually short intertrial intervals (5 min or less). Thus, the mnemonic demands imposed by these delays may have been insufficient to resolve group differences in object discrimination. It is more difficult to speculate about the potential source(s) of outcome differences in two studies where PD was modeled in rats using unilateral medial forebrain bundle (MFB) lesions similar to those shown in other studies to interfere with NOP performance as both used more standard delays (90 min, 24 hrs [139, 140]). However, in both studies rats were tested on three or four different object recognition paradigms. Thus, it is possible that some sort of practice or habituation effect associated with repeated testing experience/exposure served to narrow a performance gap that might otherwise be expected between lesioned and un-lesioned subjects. Importantly, the testing sequences used in both studies included the OiP task[139, 140]. To our knowledge these are the only other studies to use this paradigm to assess cognition and memory in a rodent model of PD. However, neither study found any negative effects of the unilateral dopamine model on OiP performance[139, 140]. Because one study used a shorter, more standard 5 min delay and the other used a longer, 24-hour delay, there are multiple experimental variables that differentiate each of the three studies where OiP has been tested in rodent models of PD. Accordingly, although it is tempting to speculate that unilateral vs. bilateral dopamine lesions are a critical difference, it is premature to identify this as causal.

### Sex Differences in Object Recognition Memory in Preclinical Rodent Models of PD

Sex differences are prominent in many features of PD including the incidence and severity of pre-dementia deficits in cognition and memory[31, 33, 37, 38]. Resolving the bases for these differences could shed light on the neural substates and pathophysiological processes that put cognition and memory at risk in PD and aid in developing better, potentially sex-specific ways to clinically manage these deficits. To date, most preclinical studies, including those examining object recognition memory noted above, have been carried out in male subjects alone. However, there are a small number of studies including the present where male and female subjects were compared. Among these, one study in rats showed that unilateral 6-OHDA MFB lesions had no impact on NOP performance in either male or female subjects[139], and one study showed that in MitoPark mice NOP discrimination was impaired in both sexes[114]. In contrast, other investigations (the present study, one in chronically reserpine treated rats[53] and one in A53T transgenic mice[52]) showed that deficits in NOP discrimination were more severe in males. Thus, similar sex differences were identified across different rodent species and using different models of PD. This suggests that factors other than these account for the variable findings for sex differences in NOP discrimination. However, one feature that consistently parses along lines of studies where sex differences were or were not observed is the duration of the intertrial interval used. Specifically, those studies showing male-specific impacts of the PD model on object recognition used shorter intertrial intervals (60-90min). However, the studies where no sex differences were reported used longer, 24hr intertrial intervals. These differences in delays map to data from lesion, transient silencing, c-fos labeling and other studies showing that perirhinal cortex is critical for NOP performance at all delays, but that the hippocampus may be more selectively engaged when inter trial intervals are longer (> 3 hrs) [141]. This provisionally identifies the perirhinal cortex as a/the critical substrate in conferring male vulnerability and female sparing of NOP performance in preclinical rodent models of PD models. This idea gains further traction from findings from this lab showing that 6-OHDA lesions sex-specifically impair performance in the OiP task (present study) but impair performance on the WWWhen ELM task equally in males and females[57]. While both tasks engage hippocampal and medial prefrontal circuits, only the OiP task is additionally linked to perirhinal cortex[99, 113]. Further, studies in humans, non-human primates and rodents link perirhinal circuits with familiarity, recency and semantic but not episodic forms of memory function[142–145]. Together, these data raise the possibility that sex differences in cognition and memory in PD reflect a vulnerability of perirhinal cortex in males and a resilience of this region in female to pathophysiological processes of disease. Although not stratified by sex, perirhinal cortical thinning has been identified in human imaging studies of PD, giving further impetus to investigate this region[146]. Findings that neostriatal 6-OHDA lesions have minimal impact on dopamine innervation in the perirhinal cortex[147] suggest that some of the non-dopaminergic, e.g., cholinergic, systems that are also at risk in PD[148] and essential for the NOP task[72, 75] should be targeted for first investigations. The present findings of sex differences in the impacts of 6-OHDA lesions on NOP and OiP but not on Barnes maze or WWWhen performance[56, 57] further suggest that sex differences in cognition and memory in this PD model are highly behavioral task- and behavioral domain-specific.

### Sex Hormone Effects on Object Recognition Memory in Preclinical Rodent Models of PD

Sex differences in PD have generated hypotheses for protective effects of ovarian steroids in females[3, 23, 25, 27, 28, 44, 45]. In the present study, investigations of hormone effects in female rats given 6-OHDA lesions were limited to comparisons of subjects that were behaviorally tested during phases of the estrous cycle when circulating ovarian steroid levels are predictably higher (proestrus or estrus) or lower (diestrus). These revealed minimal impacts of naturally occurring, low amplitude cyclical changes in ovarian steroids on NOP or OiP performance in control or 6-OHDA-lesioned rats. However, these findings are statistically underpowered and cannot be considered definitive. Rather, although not entirely consistent, clinical and preclinical studies generally indicate that elevated circulating ovarian steroid levels—both endogenous and associated with hormone augmentation have the potential to minimize disease risk, delay disease onset and reduce severity of motor and non-motor symptoms in females[21, 27, 28, 45, 47, 54, 62]. These data, along with the well-known impacts of ovariectomy and hormone replacement on object recognition memory function in rodents[66, 105, 109, 149] underscore the need for future studies to rigorously investigate the effects of sex hormones in PD models on cognition and memory in female subjects.

Compared to females, clinical studies exploring functional links between circulating hormones and PD-related risk in males are comparatively few and their outcomes to date more variable[43]. For example, some but not all studies have shown that testosterone levels are lower than normal in male PD patients[41, 42, 150, 151]. Clinical trials examining the effects of testosterone replacement on mood, motor function, cognition and memory have also yielded mixed results[46, 150, 152]. In contrast, preclinical studies in dopamine-lesioned rodents have more consistently identified correlations between the presence of circulating androgens and more severe motor or cognitive deficits, and the amelioration of these deficits in subjects where androgens are depleted[56, 58, 60]. For cognition and memory, these data come from Barnes and WWWhen testing[56, 57]. Previous studies in non-lesioned animals identified discrete measures of behavior in each task as androgen-and/or estrogen-sensitive[103, 104]. While all elements of performance were impaired by 6-OHDA lesions, only the androgen-sensitive behaviors were spared in 6-OHDA-lesioned male rats where androgens were depleted, i.e., in GDX-OH and GDXE-OH but not MALE-OH or GDXTP-OH subjects. However, in extending investigations to object recognition paradigms, different patterns of hormone sensitivity emerged. For the NOP task, where performance has been previously shown to be impaired by GDX in an androgen-sensitive, estrogen-insensitive manner[76] 6-OHDA lesions impaired discrimination to similar, significant degrees in all male groups regardless of hormone status i.e., similarly in MALE-OH, GDX-OH, GDXE-OH and GDXTP-OH rats alike. This suggests that male sex per se, i.e., organizational but not activational gonadal steroid effects, confer PD-related risk for NOP performance. These findings further indicate that in this model some but not all hormone-sensitive elements of cognition or memory can be modulated by changes in the hormonal milieu and some but not all androgen-sensitive behaviors respond favorably to androgen depletion. Findings obtained with respect to the OiP paradigm brought additional levels of specificity and complexity. First, studies in non-lesioned rats showed that GDX induced significant deficits in OiP discrimination that were attenuated in GDXE but only marginally so in GDXTP rats; this pattern implicates estrogen stimulation over and above that derived from testosterone metabolism as required to support task performance. Six-hydroxydopamine lesions were also shown to impair OiP performance in MALE-OH, GDX-OH and GDXTP-OH rats but not in the GDXE-OH group. This suggests that 6-OHDA-induced deficits in OiP in male rats are also attenuated by exogenous rather/more than endogenous estrogens. This sourcing appears to be significant as no such sparing was seen in 6-OHDA-lesioned animals for estrogen-sensitive WWWhen behaviors—which are estrogen sensitive behaviors for which active hormone can be fully accounted for by testosterone metabolism[57]. This suggests that the benefits of estrogen replacement in males – at least in this 6-OHDA lesion PD model, may be limited to those aspects of cognition or memory that are sensitive to exogenous more so or rather than endogenous estrogen stimulation. Given the very different effects that the central vs. peripheral estrogens have been shown to exert over the nigrostriatal dopamine axis[153], it will be important for future studies to rigorously evaluate cognition and memory in rodent models of PD where selective aromatase or 5-α-reductase inhibitors are used to manipulate testosterone’s metabolic fates.

### Summary and Conclusions

The data presented here and previously make a collective case for biological sex and for organizational and activational effects of sex hormones as influencing cognition and memory in a 6-OHDA lesion rat model of PD in complex behavioral task- and behavioral domain-specific ways. Findings to date show that only certain deficits that are induced by this PD model show male over female sex differences. Further, findings in males show that only some behavioral deficits induced by 6-OHDA lesions are sensitive to organizational but not activational hormone stimulation. Finally, as for protective actions, our data indicate that only some androgen-sensitive behaviors that are disrupted by 6-OHDA lesions benefit from reduced androgen signaling, and that benefits of estrogen replacement in males may be limited to lesion-induced deficits in behavioral domains that are supported more by exogenous rather than endogenous sources of estrogen. The degrees of specificity that are unfolding were unexpected and could explain why a cohesive clinical picture of endocrine impacts on higher order function in PD has remained elusive. It also seems intuitive that the differences in sex and sex-hormone sensitivities seen across tasks will map in systematic ways to the neural underpinnings that each engages. The rich extant literature describing the neural networks and neurochemical systems associated with these tasks[69-72, 74, 75] should be instructive in framing testable hypotheses as to where relevant and potentially therapeutic estrogen and androgen impacts may be levied. Although further study is needed, the distinct roles that have been established for prefrontal vs. perirhinal cortices, for discrete hippocampal subfields and for medial vs. lateral entorhinal cortices in and across different object memory-based tasks[70–72, 74] suggests that these loci may also be important in shaping their differential sensitivities to biological sex and sex hormones. Identifying these pivotal regions, i.e., determining where critical hormone effects are levied are necessary first steps for identifying how relevant hormone actions manifest. As the data in hand suggest, this could be important in addressing unmet needs for safer, more effective ways for treating cognitive and memory impairments in PD.

## ACKNOWLEDGMENTS

The authors thank Keertana Madhira for assistance in behavioral scoring, Lauren Senia for assistance with histology and Meagan Conner for assistance with behavioral testing. This work was supported by the National Institutes of Health [R21 NS11000] to MFK and by a grant from the Thomas Hartman Center for Parkinson’s Research to MFK

